# Defining the dynamic chromatin landscape of nephron progenitors

**DOI:** 10.1101/515429

**Authors:** Sylvia Hilliard, Renfang Song, Hongbing Liu, Chao-hui Chen, Yuwen Li, Melody Baddoo, Erik Flemington, Alanna Wanek, Jay Kolls, Zubaida Saifudeen, Samir S. El-Dahr

## Abstract

Six2^+^ cap mesenchyme cells, also called nephrons progenitor cells (NPC), are precursors of all epithelial cell types of the nephron, the filtering unit of the kidney. Current evidence indicates that perinatal “old” NPC have a greater tendency to exit the progenitor niche and differentiate into nascent nephrons than their embryonic “young” counterpart. Understanding the underpinnings of NPC aging may offer insights to rejuvenate old NPC and expand the progenitor pool. Here, we compared the chromatin landscape of young and old NPC and found common features reflecting their shared lineage but also intrinsic differences in chromatin accessibility and enhancer landscape supporting the view that old NPC are epigenetically poised for differentiation. Annotation of open chromatin regions and active enhancers uncovered the transcription factor Bach2 as a potential link between the pro-renewal MAPK/AP1 and pro-differentiation Six2/b-catenin pathways that might be of critical importance in regulation of NPC fate. Our data provide the first glimpse of the dynamic chromatin landscape of NPC and serve as a platform for future studies of the impact of genetic or environmental perturbations on the epigenome of NPC.

**Summary statement:** Hilliard et al. investigated the chromatin landscape of native Six2^+^ nephron progenitors across their lifespan. They identified age-dependent changes in accessible chromatin and regulatory regions supporting the view that old nephron progenitors are epigenetically poised for differentiation.

## INTRODUCTION

Reciprocal interactions between the ureteric bud and surrounding NPC of the cap mesenchyme govern nephron induction. The cap mesenchyme is composed of an early progenitor Cited1^+^/Six2^+^ compartment and a transit Cited1^−^/Six2^+^ compartment that subsequently differentiate into the pretubular aggregate, the precursor of the renal vesicle, the earliest epithelial precursor of the nephron (Brown et al., 2013; Mugford et al., 2009). Careful morphometric studies and cell cycle analyses have shown that the proportion of NPC progressing through the cell cycle decreases with NPC aging, whereas the contribution of cell death is minimal, suggesting that all NPC exit occurs via differentiation into early nephrons (Short et al., 2014). Using genetic and primary cell culture models, Fgf9 and Bmp7 were shown to stimulate the MAPK pathway activation of Fos and Jun in the cap mesenchyme leading to the formation of the AP-1 heterodimer which stimulates cell cycle and growth factor genes contributing to the maintenance of the NPC population (Muthukrishnan et al., 2015). Although Fgf9 levels do not fall appreciably during NPC aging, Fgf20 levels do (Barak et al., 2012). It is therefore possible that reduced growth factor availability/activity in the niche is partly responsible for the short lifespan of NPC. However, there are also intrinsic differences between young and old NPC. For example, Six2 (and other stemness factors such as Wt1, Osr1 and Sall1) levels decline in postnatal NPC suggesting that these low levels cannot sustain NPC stemness in the face of elevated canonical Wnt signaling. A decline in the glycolytic capacity has also been shown by RNA-seq on postnatal NPC (Chen et al., 2015) as well as in primary young and old NPC (Liu et al., 2017). These changes translate into differences in cell behavior as demonstrated in the heterochronic transplantation studies (Chen et al., 2015): whereas young NPC tend to remain in the progenitor niche, old NPC exit and differentiate. The biological underpinnings of NPC aging, i.e., the greater tendency of perinatal NPC to differentiate compared to their embryonic counterpart, are not well understood.

Here, we compared the chromatin landscape of young and old NPC and find that dynamic chromatin accessibility to developmental enhancers is an intrinsic property of aging NPC. Genome-wide ATAC-seq and ChIP-seq uncovered common and differentially accessible chromatin regions in young vs. old NPC reflecting their shared identity but also their maturational differences. Relative gain and loss of enhancer accessibility correlated with NPC gene expression and identified the poised epigenetic state of differentiation genes. While the open chromatin of young NPC is enriched in binding sites for the core NPC transcription factors (Six2, Wt1, Hoxa/c/d, Tead, AP1), old NPC gain chromatin accessibility to the Bach2/Batf complex, a repressor of AP1-mediated transcriptional activation. Importantly, Bach2 is a component of the transcription factor code of the renal vesicle, the earliest precursor of the nephron, and is a genomic target of the Six2/b-catenin complex, whereas MAPK/AP1 is required to maintain the progenitor state of NPC. In summary, our data support the notion that dynamic changes in the NPC epigenome over their lifespan balance NPC proliferation and differentiation. We propose Bach2 as a potential molecular link between the MAPK/AP1 and Six2/b-catenin pathways.

## RESULTS

### Experimental protocol (Fig. 1)

To compare the open chromatin landscape of young (embryonic) and old (perinatal) NPC, we applied the assay for transposase-accessible chromatin by high-throughput sequencing (ATAC-seq) to fluorescence-activated cell sorted (FACS) GFP^+^ cells isolated from embryonic (E13, E16) and perinatal (P0, P2) Six2^GC^ mice (Kobayashi et al., 2008), which specifically express GFP under the Six2 regulatory elements in the cap mesenchyme. By gating fluorescence and forward scatter, we also obtained enriched populations of GFP^high^ and GFP^low^ NPC from newborn mice (referred to as P0-H and P0-L), representing undifferentiated and differentiating NPC, respectively. To reduce background, remove mitochondria from the transposition reaction, and increase the complexity of the library, we used the Omni-ATAC protocol (Corces et al., 2017) on 50,000 FACS-GFP^+^ (n=3-4 biological replicates per age group). In another set of experiments, we applied magnetic-activated cell sorting (MACS) (Brown et al., 2013) to isolate the NPC from E13 and P0 CD-1 mice. To enrich for the self-renewing NPC population, E13 and P0 MACS-NPC were expanded in nephron progenitor expansion medium (NPEM) (Brown et al., 2013; Brown et al., 2015) for two passages then subjected to chromatin immunoprecipitation.

**Figure 1.**
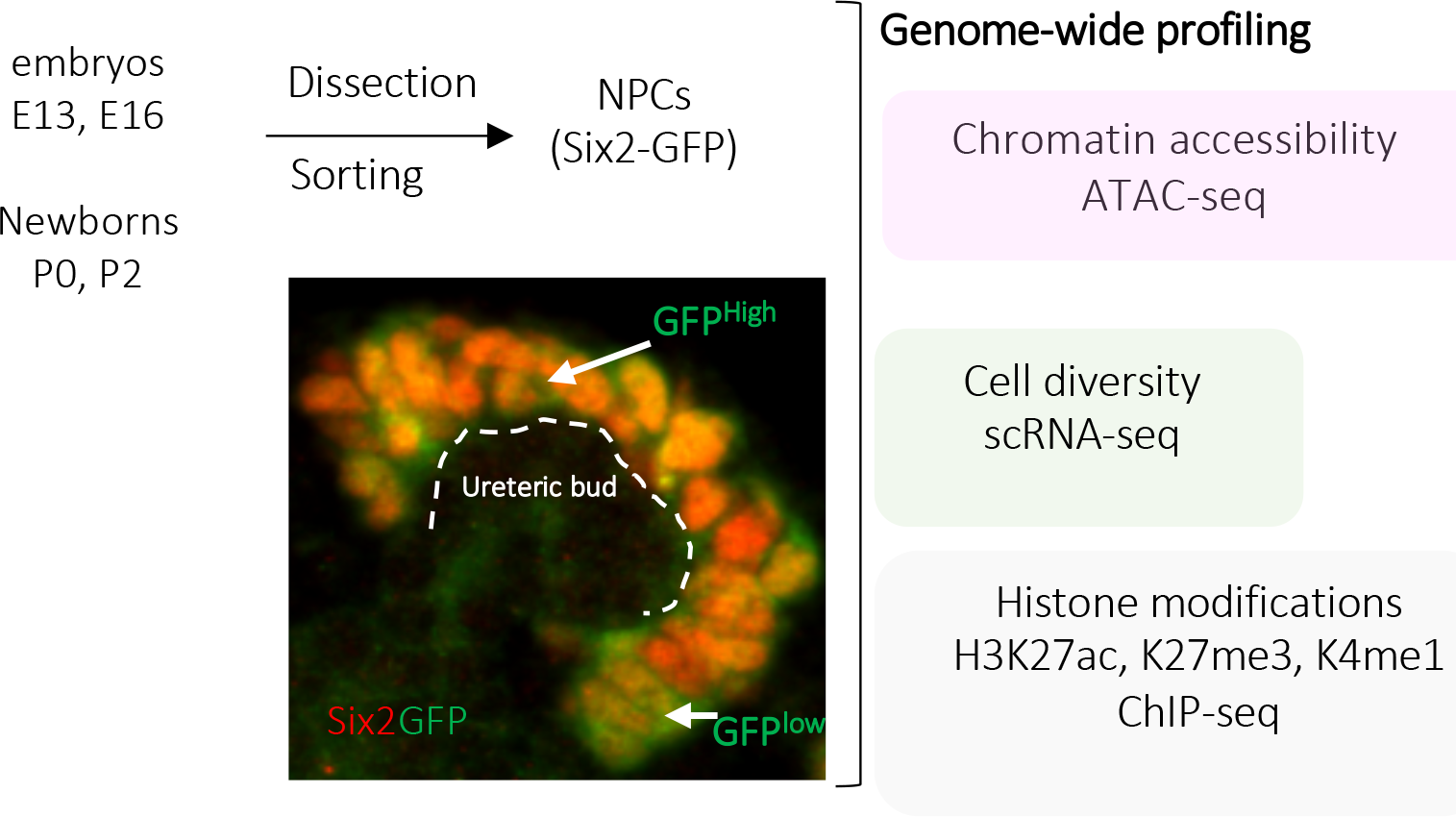
Experimental strategy for epigenetic and transcriptional profiling of nephron progenitor cells (NPC)

### Assessment of NPC diversity

We applied droplet-based single-cell RNA sequencing to 10,524 GFP^+^ cells isolated from E16 Six2^GC^ mice. Clustering analysis identified 13 distinct cell clusters varying from as few as 150 cells to as many as 1616 cells per cluster (Fig. 2A). Cluster 2 and Clusters 6-8 were enriched in NPC expressing centrosome duplication, cell cycle and DNA replication genes (Fig. 2B). Cluster 12 contained fewer numbers of NPC (343 cells, 3%) and expressed early differentiation genes such as *Wnt4, Pax8, Wfdc2 and Kdm2b* (Fig. 2C). This was not surprising since lower levels of Six2(GFP) continue to be present in early nascent elements such as the pretubular aggregate and renal vesicle. As discussed later in the text (Fig 7A), chromatin profiling of E13 and P0 NPC confirmed that the promoters of these early differentiation genes are bivalent (H3K4me1/H3K27me3) and therefore in a poised transcriptional state expressing low transcript levels. Cluster 13 is composed of NPC co-expressing stromal mesenchymal markers such as Lgals1, Maged2, and Gpc3 (Fig. 2D). A previously published study using scRNA-seq showed that individual NPC exhibit stochastic expression of stroma markers (Brunskill et al., 2014). These findings confirm that Six2^+^ NPC are composed of diverse cell populations some which are actively engaged in self-renewal while others are poised to differentiate.

**Figure 2.**
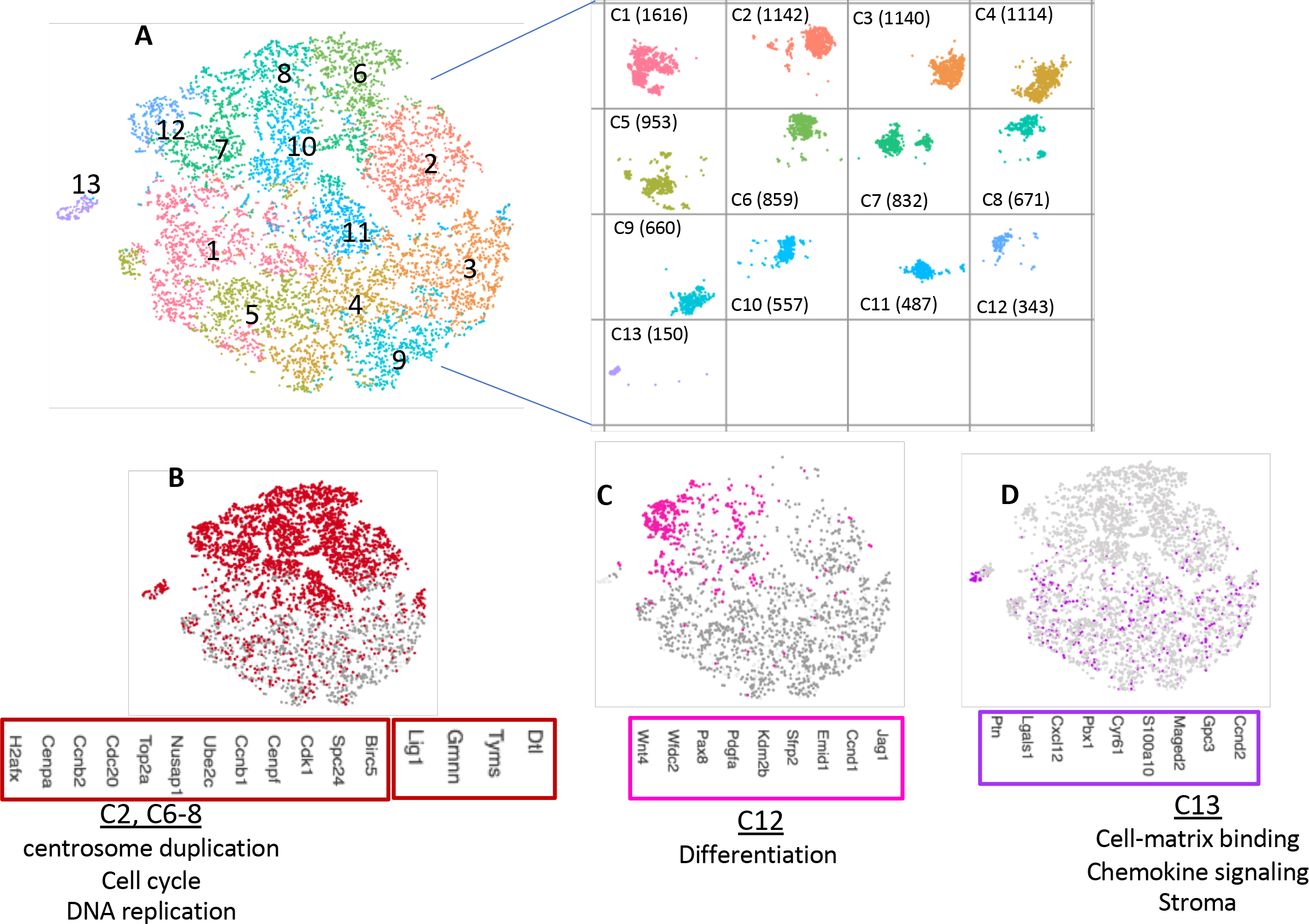
Single cell RNA-seq reveals NPC diversity. NPC from E16 *Six2GFPCre* mice were isolated by FACS and subjected to droplet-based scRNA-seq (10X Genomics platform). (A) Unbiased clustering of GFP-positive Six2-lineage NPC analyzed by singlecell RNA-seq. (B-D) Gene expression plots for marker genes; cluster numbers as indicated.

### The open chromatin landscape of NPC

ATAC-seq results were highly reproducible between biological replicates and showed a clear enrichment at the regulatory elements. As an example (Fig. 3A), in the *Six2* locus, the replicates show that the open chromatin regions are concentrated in the promoter region and the annotated enhancer located 60kb upstream of the TSS (Park et al., 2012). Analysis of ATAC-seq peaks revealed that young NPC possessed more distal open chromatin regions than older NPC (Fig. 3B). Heatmap clustering of ATAC peaks in annotated genes around the TSS (−1 to +1 kb) did not reveal significant differences in open chromatin in E16 and P2 NPC (Fig. 3C). Gene Ontology of the distal (−1 to −60 kb) regions showed common and distinct gene signature sets and pathways associated with the open chromatin regions in E16 and P2 NPC (Fig. 3D). Among the common features are regulation of stem cell maintenance and nephrogenesis as well as Notch signaling and the TCA cycle. Distinct functional signatures include response to reactive oxygen species and pentose phosphate pathway (E16 NPC) and regulation of cell shape and ATP synthesis (P2 NPC).

**Figure 3.**
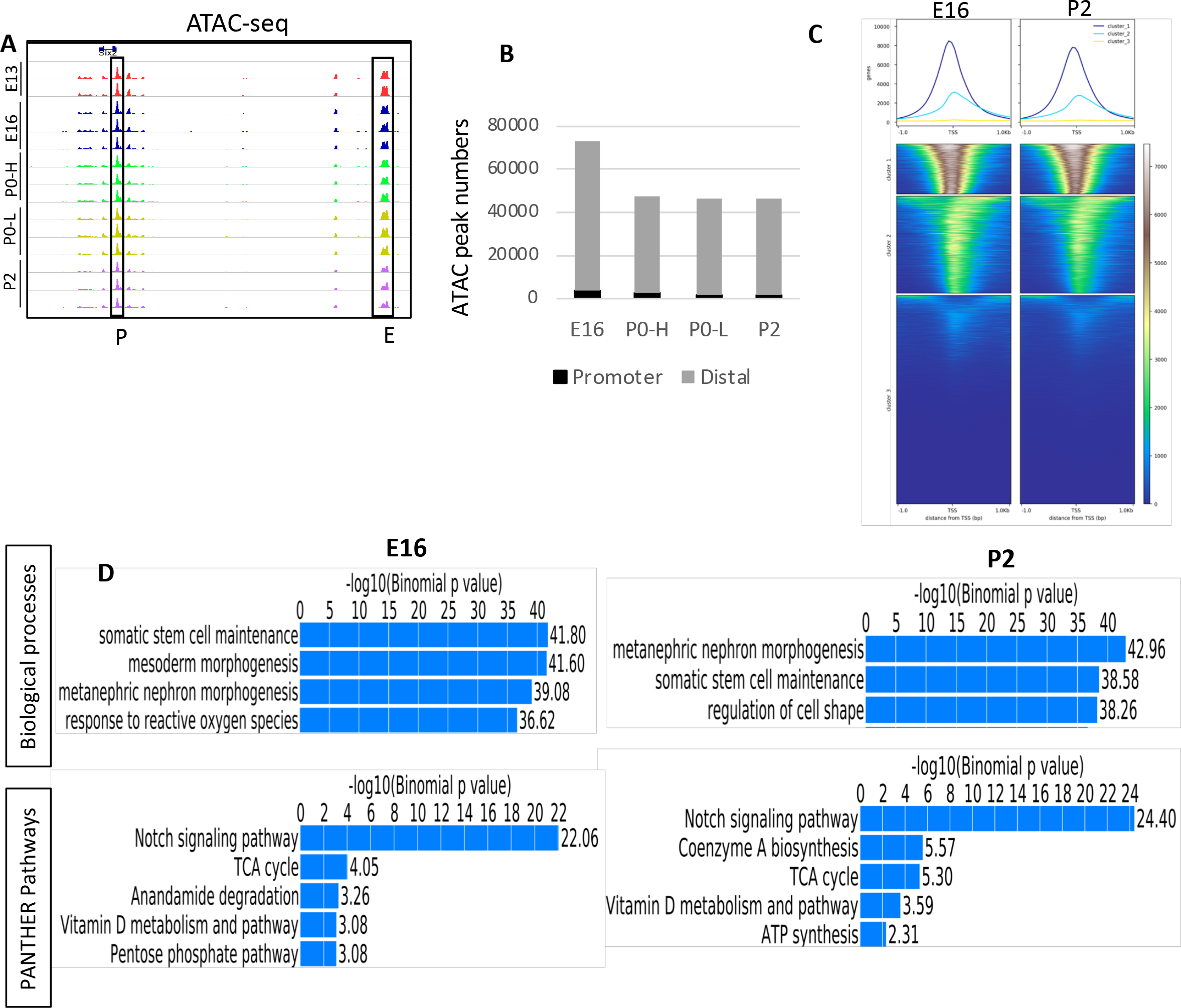
Profiling of open chromatin regions in young and old NPC by ATAC-seq reveals common and distinct biological features. (A) Representative ATAC-seq profiles of biological replicates from young (E13, E16) and old (P0, P2) NPC. P0-H and P0-L represent GFP^high^ and GFP^low^ perinatal NPC. (B) Total numbers of ATAC-seq peaks separated into promoter (±1 kb transcription start site) and distal genomic regions (−1 to −60 kb) in NPC of various ages. (C) Heatmap of ATAC peaks in the −1 to +1 kb around TSS in annotated genes at E16 and P2 NPC. (D) Functional annotation of ATAC peaks by GREAT analysis in young and old NPC.

### NPC aging is associated with differential chromatin accessibility

We compared the open chromatin regions of E16 and P2 NPC as well as of P0-H and P0-L NPC using DiffBind R (http://bioconductor.org/packages/release/bioc/vignettes/DiffBind/inst/doc/DiffBind.pdf). The affinity analysis is a quantitative approach to assess for differential chromatin access at consensus peaks. This method takes read densities computed over consensus peak regions and provides a statistical estimate of the difference in read concentration between the two conditions. Heatmap correlation plots, Principal Component Analysis, and MA plots highlight the differences in ATAC read concentrations between E16 and P2 NPC and P0-H and P0-L NPC (Fig. 4 A-C, F-H). The MA plots, shown in Fig. 4C,H depict the differences between ATAC signals in the two experimental samples by transforming the data onto M (log ratio) and A (mean average) scales, then plotting these values. Differential chromatin accessibility depicted in the MA plots is expressed as a log fold change of at least 2-fold and a p-value of < 0.05. The MA plot in Fig. 4C shows the relative gain of open chromatin regions in E16 NPC (above the 0 threshold line) as compared to the gain in P2 NPC (below the 0 threshold line). The MA plot in Fig. 4H depicts the relative gain of open chromatin regions in P0-H (above the 0-threshold line) as compared to the gain in P0-L NPC (below the 0-threshold line). NPC aging is associated with a decline in differential chromatin accessibility of the stemness factor, *Six2*, but a reciprocal increase in chromatin accessibility to the differentiation transcription factor *HNF1-b* (Fig. 4 D,E). Likewise, differentiating P0-L NPC gain chromatin accessibility in differentiation genes (e.g., Lef1, Irx, Hnf1b, Tcf7l2 and some of the Notch components) (Fig. 4 I). P0-L NPC also displayed increased accessibility around the TSS as compared to P0-H NPC (Fig. 4 J). Collectively, these findings illustrate the dynamic open chromatin landscape of NPC during maturation and are consistent with the view that old NPC are epigenetically programmed to differentiate.

**Figure 4.**
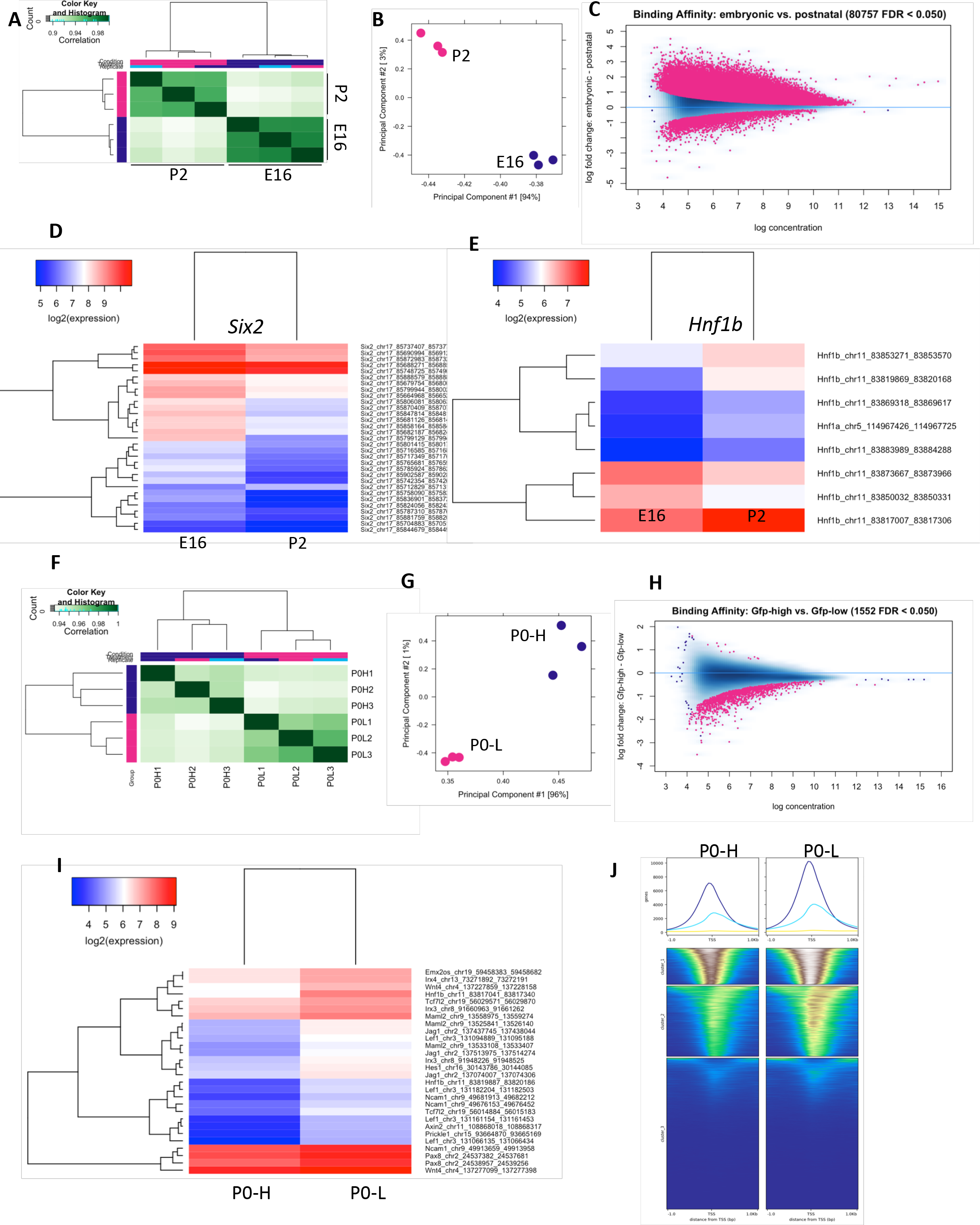
Differential chromatin accessibility in young and old NPC. Heatmap (A,F), Principal Component Analysis (B, G) and MA plot (C,H) of ATAC-seq peaks in E16 vs. P2 NPC or P0-H vs. P0-L NPC. (D,E) Differential chromatin accessibility at the *Six2* and HNF1b locus between E16 vs. P2 NPC. (I) Differential chromatin accessibility upon differentiation between P0-H and P0-L NPC. (J) Heatmap of ATAC peaks in the −1 to +1 kb around TSS in annotated genes at E16 and P2 NPC.

### ATAC peaks correlate with NPC gene expression and transcription factor binding

We also compared our ATAC-seq signals representing open promoters and enhancers with published gene expression of NPC lineage (O’Brien et al., 2018). The results showed that progenitor genes in E16 NPC (e.g., Six2, Osr1 and Meox1) exhibit more open chromatin at distal elements and promoters than P2 NPC (Fig. 5A). In comparison, poised genes in P2 NPC (e.g., Wnt4, Sulf1 and Mafb) exhibit more open chromatin features than E16 NPC (Fig. 5A). These findings indicate a good correlation between chromatin accessibility and gene expression levels during NPC aging.

**Figure 5.**
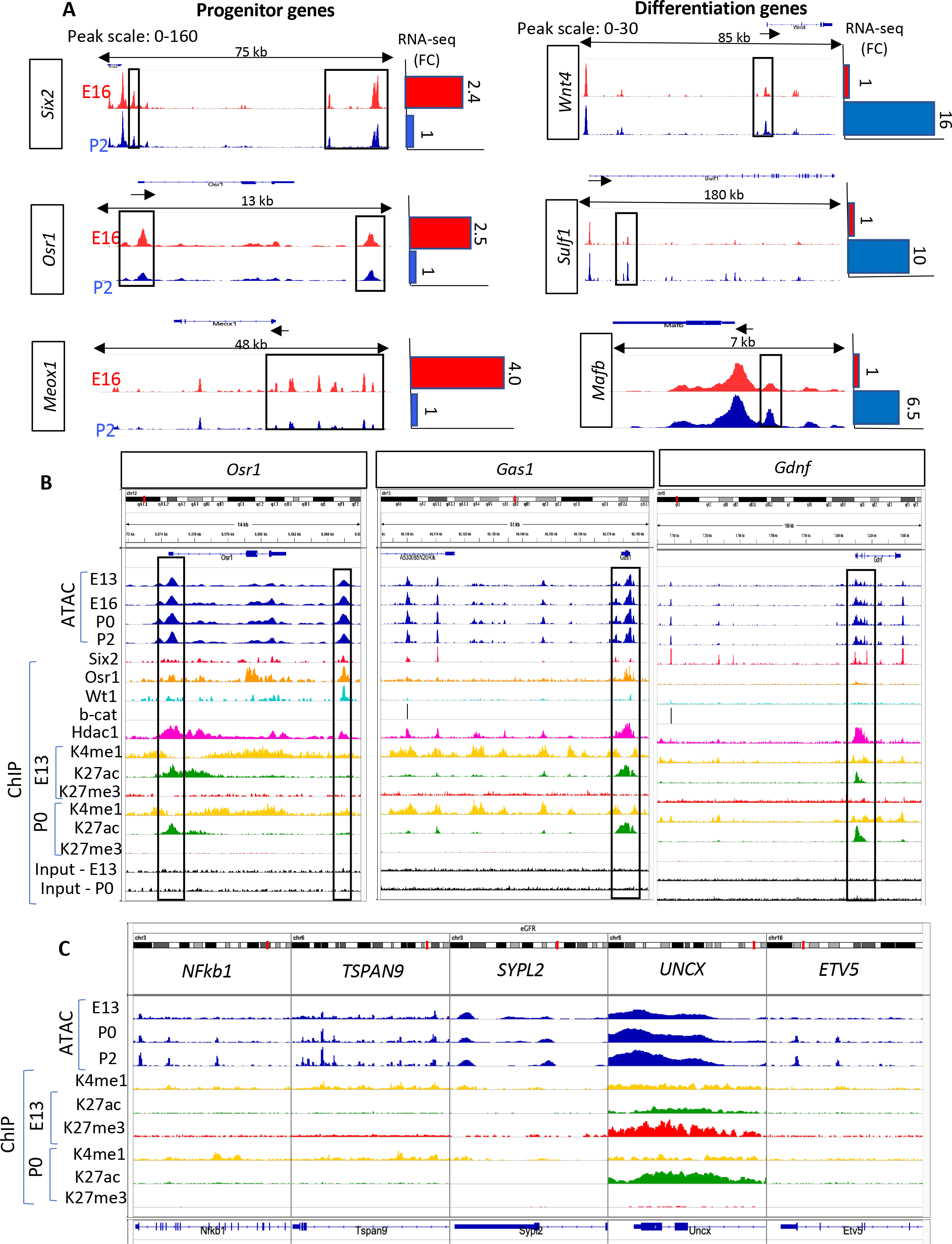
Open chromatin profiles correlate with gene expression and transcription factor binding at annotated and putative enhancers of NPC developmental regulators. (A) ATAC-seq and RNA-seq of progenitor and differentiation genes at E16.5 and P2 NPC. (B) IGV tracks showing integration of ATAC-seq and ChIP-seq peaks in promoters and enhancers of the progenitor genes, *Osr1, Gas1* and *Gdnf*. (C) ATAC/ChIP signature of genes found in genome-wide association studies of estimated glomerular filtration rate to demonstrate preferential mapping of associated variants to regulatory regions in kidney but not extra-renal tissues (Pattaro et al., 2016).

The open chromatin regions indicated by the ATAC peaks also correlated with the ChIP-seq peaks representing active enhancers (H3K4me1/H3K27ac) — this study – as well as with binding sites of the NPC core transcription factors Six2, Osr1 and Wt1 – previously shown by O’Brien et al. (O’Brien et al., 2018) (Fig. 5B). Of interest, we found that the binding of Hdac1 was a reliable indicator of open chromatin at the promoter region of actively transcribed genes where rapid cycles of histone acetylation and deacetylation are required for resetting the chromatin of active genes (Wang et al., 2009) (Fig. 5B).

### NPC chromatin profiles in genes implicated in renal function

We examined the ATAC signature of genes found in genome-wide association studies of estimated glomerular filtration rate and demonstrate preferential mapping of variants to regulatory regions in kidney but not extra-renal tissues (Pattaro et al., 2016). Although many of these genes were not active in NPC, they displayed open chromatin regions and were marked by H3K4me1 on putative primed enhancers (Fig. 5C). The homeobox transcription factor, *UNCX*, showed broad open chromatin regions that increase in width with NPC aging and is bivalent in E16 NPC but active in P2 NPC (Fig. 5C).

### The histone modifications landscape of pro-renewal and differentiation pathways

Given that the niche microenvironment (growth factor availability) may not be similar in native young vs. old NPC, we examined the epigenetic states of MACS-isolated E13 and P0 NPC grown in NPEM for two passages. The MACS isolation, which is performed on wild-type tissue, also avoids the Six2GFPCre transgene, in case it had a non-specific effect on the chromatin landscape. ChIP-seq analysis was then performed to map active (H3K4me1/H3K27ac), repressed (H3K27me3) and poised (H3K4me1/H3K27me3) enhancers. Fig S1 A-D depicts that pro-renewal pathway genes (Akt signaling, cell cycle, RTK signaling, and epigenetic regulators) are all decorated with active histone marks, the exception being cell cycle inhibitors which are occupied by broad regions of the repressive mark H3K27me3 (Fig S1B). In comparison, Fig. 6A shows that genes required for mesenchyme-to-epithelium transition are occupied by bivalent chromatin regions (H3K4me1/K27me3) indicating they are silent but poised for transcription. Furthermore, poised differentiation genes showed a decrease in occupancy in the repressive H3K27me3 mark and a reciprocal gain of H3K27ac with NPC aging (Fig. 6 A,B). Finally, non-lineage genes, e.g., pro-neural developmental genes, are occupied by broad domains of H3K27me3 which serve to restrain their expression (Fig. 6C).

**Figure 6.**
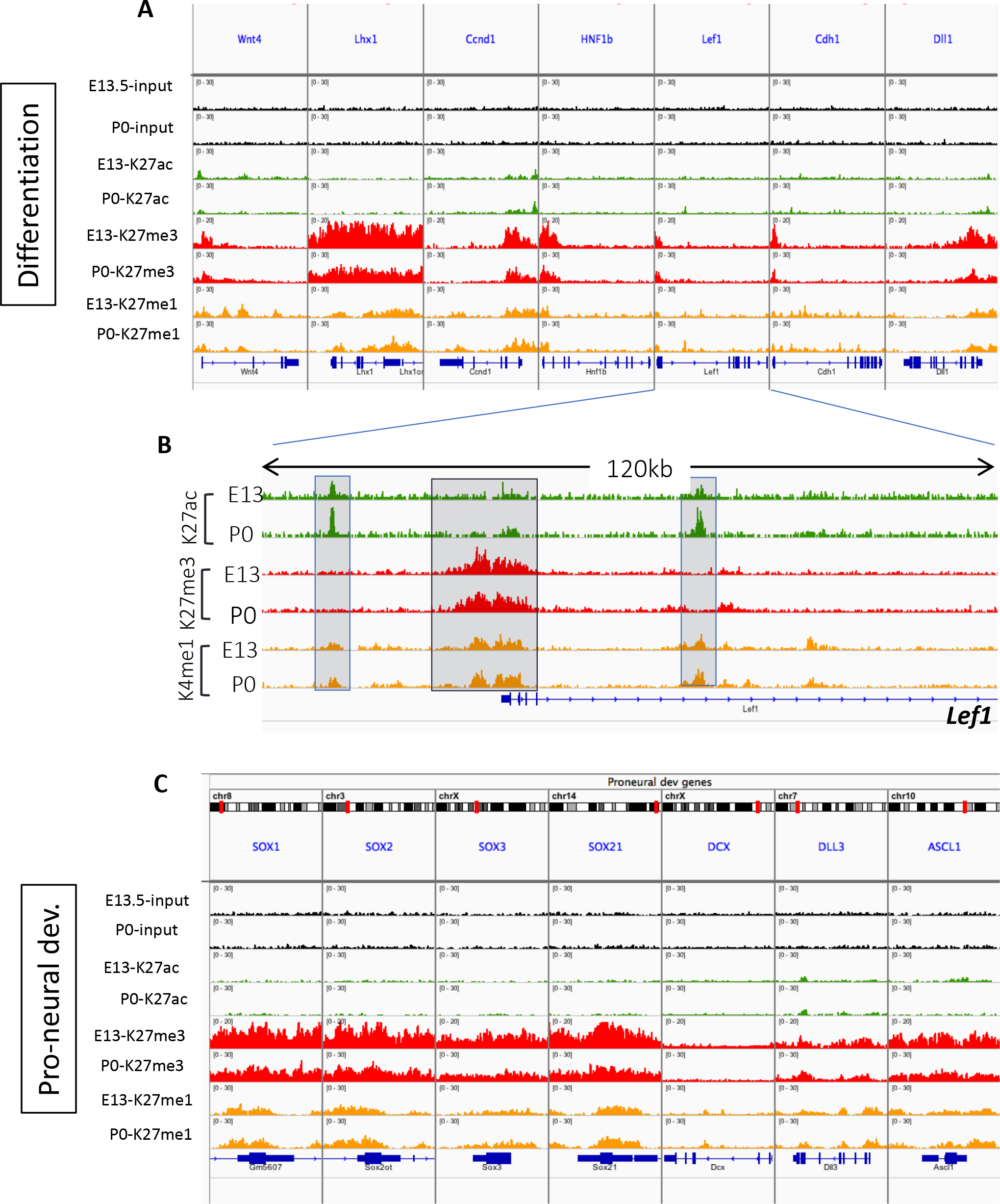
ChlP-seq profiling of histone modifications in poised differentiation (A,B) and non-lineage proneural (C) genes in young and old NPC.

Collectively, these results indicate that young and old NPC exhibit intrinsic differences in chromatin accessibility and histone modifications at annotated and putative promoters and enhancers.

### Chromatin profiling links the transcription factor Bach2 with NPC aging

We used ATAC-seq to identify transcription factor motifs within the accessible distal regions in young and old NPC. HOMER revealed enrichment of the DNA-binding motifs of the master NPC transcription factors Six2 and Wt1 across all age groups reflecting their shared lineage (Table 1 and Fig S2). Hoxc9 and Tead DNA-binding motifs were also enriched among the top 21 most enriched motifs. Hoxa11 and Hoxd11 motifs, on the other hand, were enriched in young E13 NPC but not older NPC. Notably, binding motifs for AP-1, BATF and especially Bach2 were progressively more enriched in older and differentiating NPC (Table 1 and Fig S2).

**Table 1.**
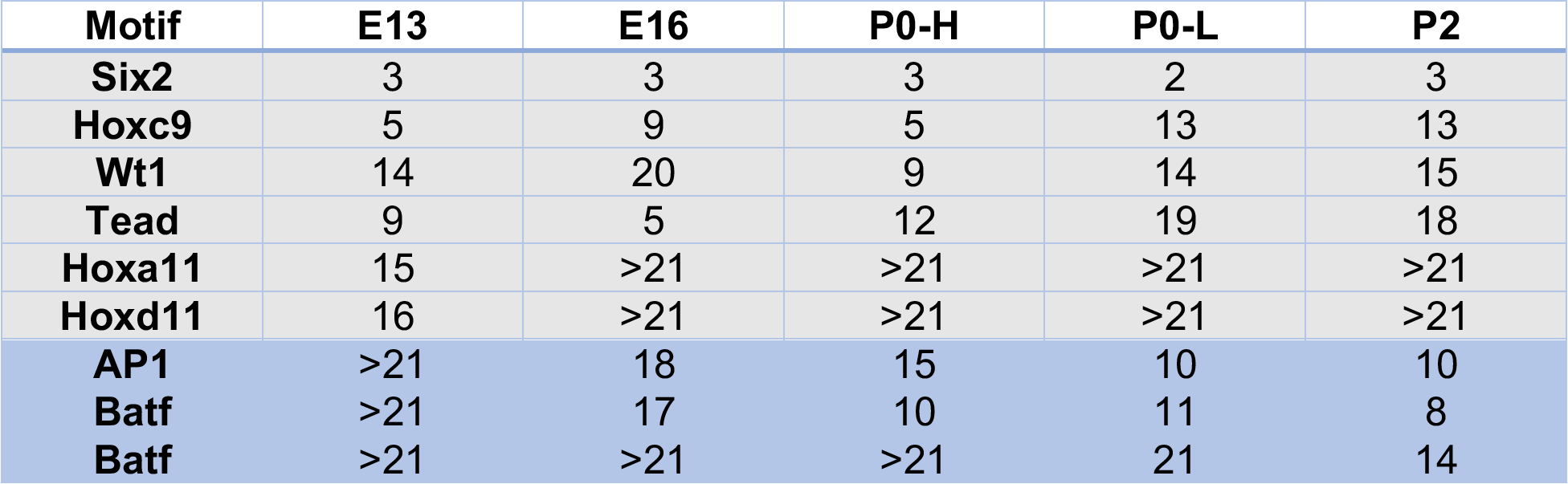
Ranking of top transcription factor motifs in ATAC-seq open chromatin peaks within the distal upstream regions (−1 to −60 kb of TSS). Rank range: 1-21.

Using ChIP-seq, we identified 18692 primed enhancers (marked by H3K4me1) at E13 compared to 27983 primed enhancers at P0 (P0>E13 +1.5-fold) (Fig. 7A). The numbers of H3K27ac-marked active enhancers were 3333 at E13 and 16207 at P0 (P0>E13 +4.8-fold) (Fig. 7B). Motif enrichment analysis of E13 NPC revealed that among the 15 most enriched active enhancers were the core factors AP1, Sox4, Hoxc9, Hoxd11 and Six2 (Fig. 7C). At P0, the 15 most enriched active enhancers were the core TFs plus Pax8, Batf, Rbpj, AP1 and Bach2 (Fig. 7D). Together, the ATAC and ChIP data highlight the age-related enrichment of Bach2/Batf motifs suggesting a temporal association between age-related changes in chromatin accessibility and the binding of AP1/Bach2/Batf.

**Figure 7.**
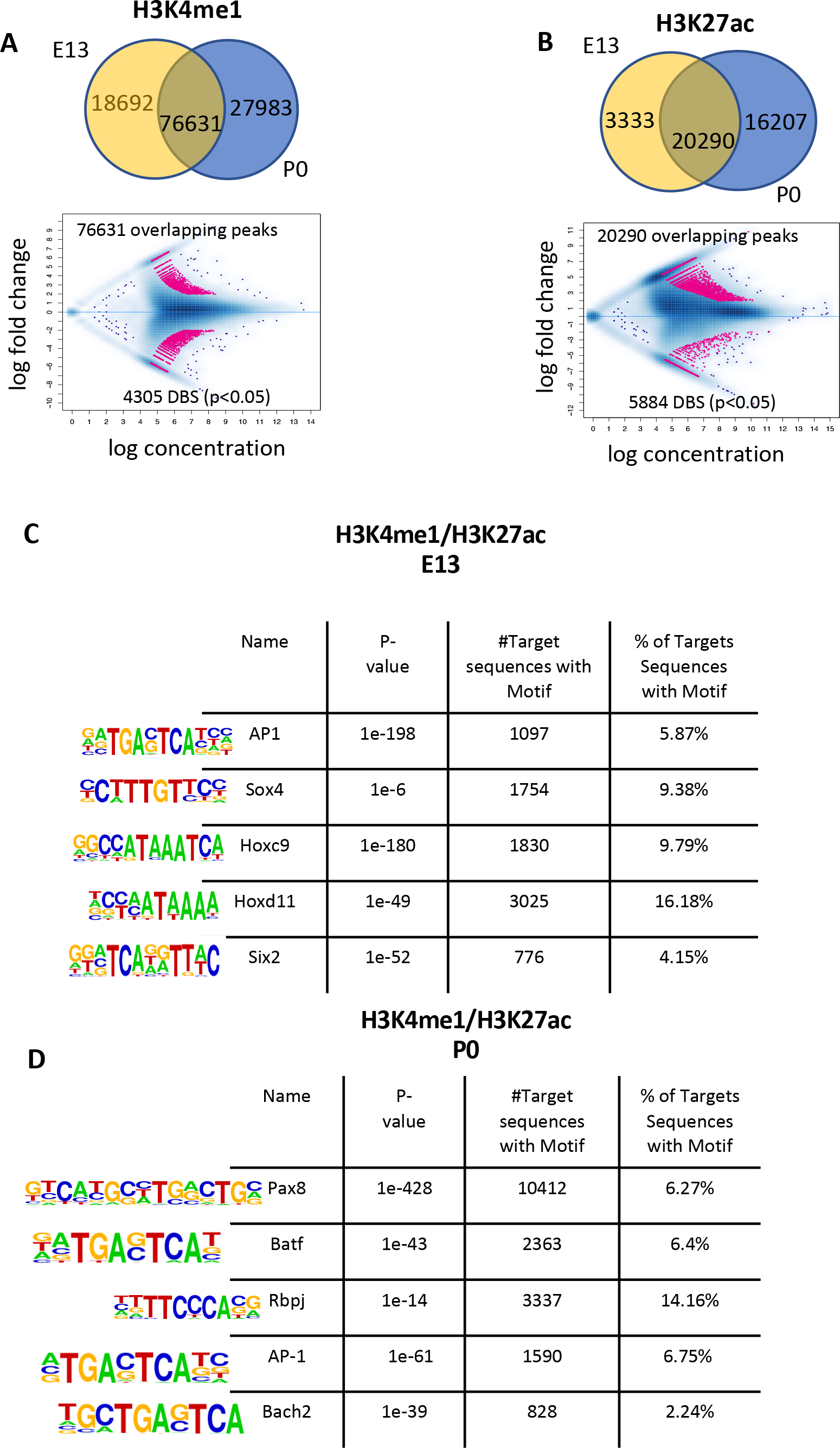
ChIP-seq of E13 and P0 NPC grown in NPC expansion medium reveals age-related gain of enhancers and confirms enrichment with AP1, Bach2 and AP1 motifs. (A,B) Venn diagrams of unique and shared H3K4me1 and H3K27ac active regions. Affinity binding analysis performed on the shared regions reveals a net gain of enhancers in P0 NPC. (C) HOMER analysis at E13 and P0 active enhancers defines distinct sets of enriched motifs in E13 and P0 NPC.

Bach2 (BTB domain and CNC Homolog 2) is a transcription factor that is enriched in immune cells and plays a key role in regulation of the developmental B-cell transcriptional programs (Zhou et al., 2016). Previous studies have shown that Bach2 acts by recruiting Batf and competes with AP-1 for sequence-specific DNA binding on target genes (Kuwahara et al., 2016; Roychoudhuri et al., 2016). Importantly, our scRNA-seq indicates that Bach2 is expressed in the NPC along with Batf and the AP1 components Junb and Atf3 (Fig. 8A). Furthermore, interrogation of the GUDMAP/RBK database revealed that *Bach2* is expressed in the distal part of the renal vesicle, the earliest nephron precursor (Fig. 8B). ATAC and ChIP-seq also showed the presence of two putative regulatory elements in the *Bach2* gene marked by the presence of consensus ATAC/H3K27ac/Six2/b-catenin peaks (Fig. 8C), suggesting that *Bach2* is a genomic target of canonical Wnt signaling, the principal driver of NPC differentiation.

**Figure 8.**
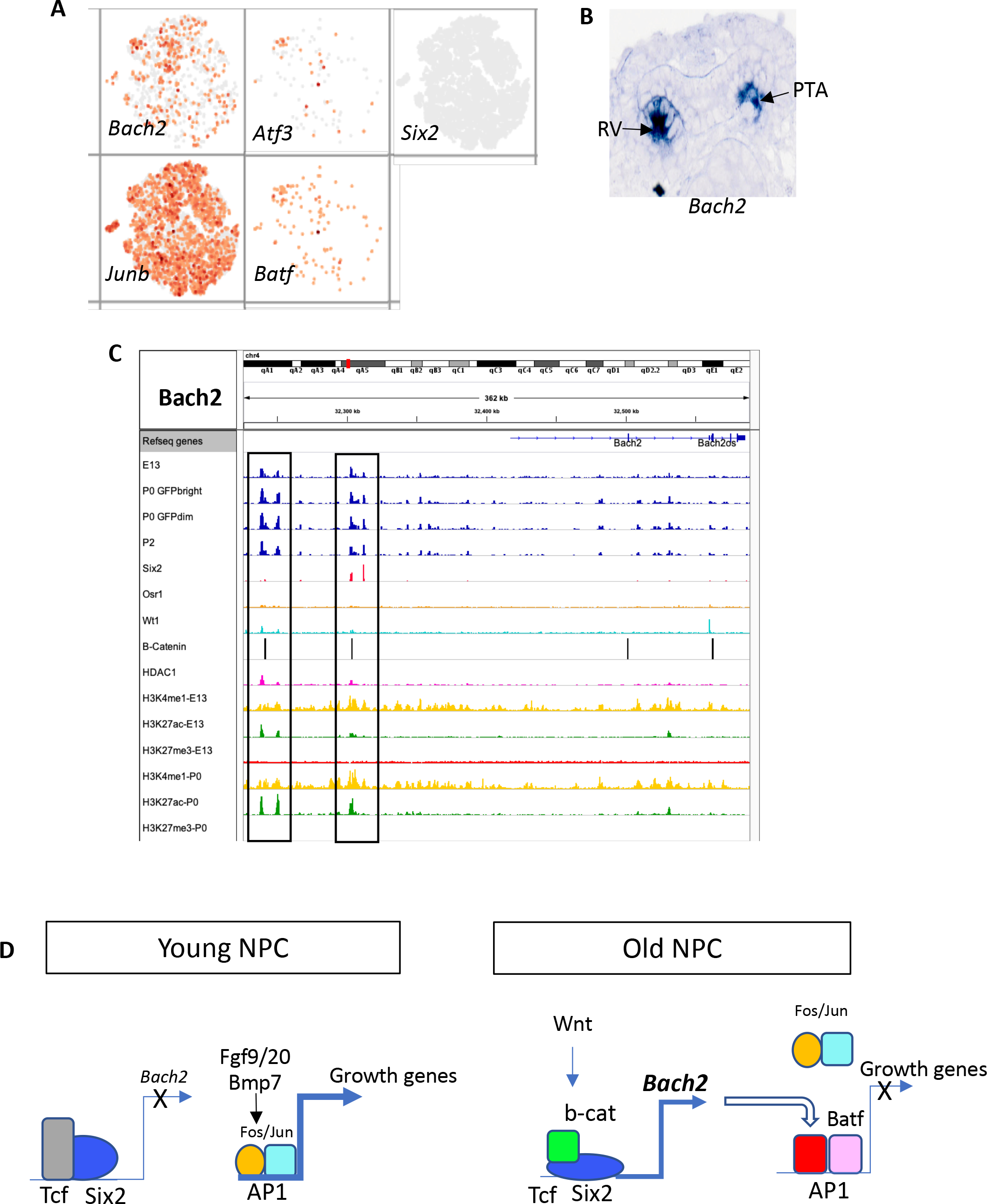
The transcription factor, *Bach2*, is expressed in the early differentiating nephron epithelium and displays active enhancers bound by Six2/β-catenin. (A) scRNA showing expression of *Bach2/Batf* transcripts and AP1 components in NPC. (B) *in situ* hybridization in TS23 stage developing mouse kidney showing that *Bach2* is expressed specifically in the pretubular aggregate (PTA) and distal renal vesicle (RV) (Source: Gudmap.org). (C) IGV tracks of the *Bach2* locus: note age-related opening of chromatin in the distal upstream region of *Bach2* (small arrow). These peaks are congruent with binding of ChIP-seq peaks for Six2 and β-catenin at two active enhancer elements (boxed regions). (D) A transcriptional model for chromatin state transition during the NPC lifespan. In young NPC, Bach2 (a differentiation-promoting gene) is repressed by Six2/Tcf, whereas genes involved in stemness maintenance are activated by the growth factor/AP1 signaling module. As the NPC ages, Six2 levels decline and canonical Wnt activity is enhanced leading to Bach2 induction. Bach2/Batf in turn competes for DNA binding with the AP1 complex to turn off expression of renewal genes.

## DISCUSSION

Understanding the epigenetic mechanisms governing the nephron progenitor life span (Adli et al., 2015) is of great translational importance, since low nephron number predisposes to hypertension and chronic kidney disease later in life. Importantly, monogenic mutations account for only about 20% of all cases of abnormal kidney development. Accordingly, there is a critical gap in our knowledge of how adverse prenatal environmental events affect renal development in the offspring. Moreover, a better understanding of the physiological, metabolic and epigenetic underpinnings of NPC aging may open new avenues to expand the nephron progenitor pool.

We previously interrogated the histone signatures of cultured metanephric mesenchyme cell lines (McLaughlin et al., 2013) and found that the onset of differentiation is accompanied by resolution of chromatin bivalency conforming to prior observations in pluripotent cells wherein genes essential for lineage commitment carry permissive histone modifications and appear poised for differentiation. In these studies, we observed that Wnt-responsive differentiation genes gain β-catenin/H3K4me3 binding and lose H3K27me3 at the TCF/LEF binding sites. By contrast, the promoters of progenitor genes show a gain of repressive marks and the loss of the active histone marks. Interestingly, immunolocalization studies in developing kidneys revealed that Six2^+^ nephron progenitors have higher levels of the repressive H3K9me2 and H3K27me3 marks and the histone modifiers, G9a, Ezh2 and HDACs (Liu et al., 2018; McLaughlin et al., 2014). While these studies were informative, additional work was needed to clarify whether the dynamic chromatin landscape observed in metanephric mesenchyme cell lines applies to native NPC.

In the present study, using native freshly isolated NPC across their lifespan, we demonstrate that as the NPC age in the niche, chromatin accessibility to the regulatory regions of poised differentiation genes increases, likely preparing these gene networks for activation of the epithelial nephrogenesis program. These changes are not simply the results of changing proportions of the self-renewing and differentiating NPC populations but likely intrinsic, since young and old NPC grown in the same growth factor expansion medium also show significant differences in their chromatin enhancer landscape. We speculate that these maturational changes in chromatin accessibility are likely orchestrated by concomitant changes in the epigenetic machinery such as the Polycomb complex, ATP-dependent chromatin remodelers, DNA methylation and the NuRD/HDAC complex, that govern the access of master regulators to their cis-acting elements. Indeed, there is genetic evidence that perturbations of the epigenetic machinery disrupt the balance between NPC proliferation and differentiation in vivo (Denner and Rauchman, 2013; El-Dahr and Saifudeen, 2018; Liu et al., 2018; Zhang et al., 2018).

A new finding of this study is that NPC aging is associated with enhanced accessibility at the Bach2/Batf occupancy sites. As mentioned earlier in the text, Bach2 was identified as part of the transcription factor signature of the distal renal vesicle, a compartment that receives high levels of Wnt9b/ß-catenin signaling (Brunskill et al., 2014). This leads us to the following working model (Fig. 8D). In young NPC, Six2/co-repressor complexes inhibit *Bach2* transcription. With further activation of Wnt/ß-catenin signaling and concomitant decline in Six2 levels, the Six2/TCF repressor is replaced by ß-catenin/TCF/co-activator complex leading to induction of *Bach2* transcription. The Bach2/Batf complex subsequently displaces the AP-1 complex inhibiting expression of AP1 targets (e.g., cell cycle genes). It is also conceivable that Bach2 targets differentiation genes to activate them. To our knowledge, studies of renal development have not been reported in Bach2- or Bach1/Bach2-deficient mice. Future investigations of *Bach2* function in nephrogenesis and delineation of Bach2-target genes will shed light on some of these gene regulatory networks. If indeed Bach2 links the proliferation and differentiation pathways in the NPC, targeting Bach2 may be a useful tool to manipulate the fate and lifespan of the NPC in renal regenerative medicine.

## MATERIALS AND METHODS

### Isolation of NPC

NPC were isolated from E13.5, E16.5, P0 and P2 CD1 mice or *Six2GFPCre* mice by magnetic-activated cell sorting (MACS) (Brown et al., 2015) or fluorescent-activated cell sorting (FACS). Animal protocols utilized in this study were approved by and in strict adherence to guidelines established by the Tulane University Institutional Animal Care and Use Committee.

### ATAC-seq

For sample library preparation we followed the Omni-ATAC method outlined by (Buenrostro et al., 2015; Corces et al., 2017). Briefly 50,000 nuclei from FACS-sorted cells were processed for Tn5 transposase-mediated tagmentation and adaptor incorporation at sites of accessible chromatin. This reaction was carried out using the Nextera DNA Library Prep kit (FC-121-1030, Illumina) at 37°C for 30min. Following tagmentation the DNA fragments were purified using the Zymo DNA Clean and Concentrator Kit (D4014, ZYMO Research). Library amplification was performed using the Ad1 and any of Ad2.1through Ad2.12 barcoded primers (Buenrostro et al., 2015). The quality of the purified DNA library was assessed on 6%TBE gels as well as on a Bioanalyzer (2100 Expert software, Agilent Technologies) using High Sensitivity DNA Chips (5067-4626, Agilent Technologies Inc.). The appropriate concentration of sample was determined using the Qubit Fluorometer (Molecular Probes). Four nM samples were pooled and run on a NextSeq 500/550 Hi Output Kit (20024907, Illumina, Inc. San Diego, CA) and the NextSeq 500 Illumina Sequencer to obtain paired end reads of 75bp. Three to four independent biological replicates were sequenced per sample.

#### Read processing and normalization of data

The paired-end reads for each sample run across 4 lanes of the flow cell (20022408, Illumina) were concatenated to obtain one forward and one reverse fastq.gz files each. The quality of the reads was assessed using FASTQC (v0.11.7). The paired end reads were aligned to the mouse reference genome mm10 using Bowtie 2. The properly aligned reads were filtered for mitochondrial reads (Sam tools) and cleared of duplicates (Picard-tools, v1.77). Only paired reads with high mapping quality (MAPQ >30) were included in the downstream analysis. The narrow peaks were called using MACS2 using the following parameters (effective genome size = 1.87e+09; - -nomodel −p 0.001 - - no lambda; band width = 300, d = 200; p value cut off = 1.00e−03). Normalized bigwig files were generated using bedtools. Annotation and Known as well as de novo Motif discovery was achieved with Hypergeometric Optimization of Motif Enrichment (HOMER). Gene ontology analysis was performed using GREAT analysis 3.0. Heatmaps, density plots and differentially mapped regions were generated using DiffBind (Bioconductor).

### Single Cell RNA-seq

We performed gene expression profiling of approximately 10,000 individual cells by using 10x Single Cell RNAseq technique provided by 10× Genomics. We first obtained a single cell suspension where cell viability was 88% or higher. These cells were applied for GEM generation and barcoding using the manufacturer’s protocol. 10×™ GemCode™ Technology allows the partition of thousands of cells into nanoliter-scale Gel Bead-ln-EMulsions (GEMs) with application of ~750,000 barcodes to separately index each cell’s transcriptome. After GEM reaction mixture, full-length barcoded cDNA were generated and amplified by PCR to generate sufficient mass for library construction. Following enzymatic fragmentation, end-repair, A-tailing, and adaptor ligation, indexed libraries were generated and sequenced. We used Cell Ranger version 2.1.1 (10×^™^ Genomics^™^) to process raw sequencing data and Seurat suite version 2.2.1 for downstream analysis. Filtering was performed to remove multiplets and broken cells and uninteresting sources of variation were regressed out. Variable genes were determined by iterative selection based on the dispersion vs. average expression of the gene. For clustering, principal-component analysis was performed for dimension reduction. Top 10 principal components (PCs) were selected by using a permutation-based test implemented in Seurat and passed to t-SNE for visualization of clusters.

### ChlP-seq

Chromatin was prepared from E13.5 and P0 NPC using the Diagenode iDeal ChIP-seq kit for Histones. Samples corresponding to 0.5 million cells were resuspended in 100 μl of shearing buffer iS1 and sheared during 8 cycles of 30 seconds “ON” / 30 seconds “OFF” with the Bioruptor Pico combined with the Bioruptor Water cooler. The shearing efficiency was analyzed using an automated capillary electrophoresis system Fragment Analyzer (High sensitivity NGS fragment kit) after RNAse treatment, reversal of the crosslinking and purification of DNA. Based on optimization conditions, we used optimal antibody quantity resulting in higher enrichment and lower background (1 μg each of anti-H3K4me1, anti-H3K27me3 and anti-H3K27ac). The antibodies used were the following: H3K27ac (Diagenode antibody, C15410196, lot A1723-0041D), H3K4me1 (Diagenode antibody, C15410194, lot A1862D), H3K27me3 (Diagenode antibody, C15410195, lot A1811-001P). H3K4me3 (Diagenode antibody, C15410003, lot 5051-001P) was used as a ChIP Positive Control. After the IP, immunoprecipitated DNA was analyzed by qPCR to evaluate the specificity of the reaction. The primers pair, Six2 Prom (Fwd3 _ Rev 4) was used as positive control for the H3K27ac mark, Mouse Negative Control Primer Set1 (Commercially available from Active Motif) was used as negative control region. Myogenic differentiation antigen 1 9MYOD1) and Mouse Negative Control Primer Set 3 were respectively used as positive and negative control regions for the H3K27me3 mark. Finally, Paired box 8 (Pax8int) and Mouse Negative Control Set 1 were respectively used as positive and negative controls for H3K4me1 mark. An IP with a control isotype (IgG 1 lμg) was also performed. 500 pg of DNA was subsequently used for library preparation using the MicroPlex v2 protocol. The ChIP samples (8 samples in total) were processed together. A control library was processed in parallel of the samples using the same amount of a control Diagenode ChIP’s DNA. According to the protocol, 12 cycles of amplification were performed to amplify the libraries. After amplification, 1 μl of each library was loaded on BioAnalyzer for quality and quantified using the Qubit ds DNA HS kit. Reference genomes were obtained from the UCSC genome browser. Sequencing was performed on an Illumina HiSeq 2500, running HiSeq Control Software 2.2.58. Quality control of sequencing reads was performed using FastQC. Reads were then aligned to the reference genome using BWA v. 0.7.5a. Samples were filtered for regions blacklisted by the ENCODE project. Subsequently samples were deduplicated using SAMtools v. 1.3.1. Alignment coordinates were converted to BED format using BEDTools v.2.17 and peak calling was performed using Sicer. The main ChIP-Seq statistics are shown in Supplemental Table S2. Peaks sets generated with peak calling analysis were analyzed using DiffBind R/Bioconductor package.

## Acknowledgments

The Authors are grateful to Kejing Song (Tulane Genomic and epigenomic and sequencing cores), and the laboratory of Dr. Mazhar Adli (University of Virginia) for help in the early phase of the study.

## Data availability

The files have been deposited in NCBI GSE124804.

---

**Supplemental Table 1:**
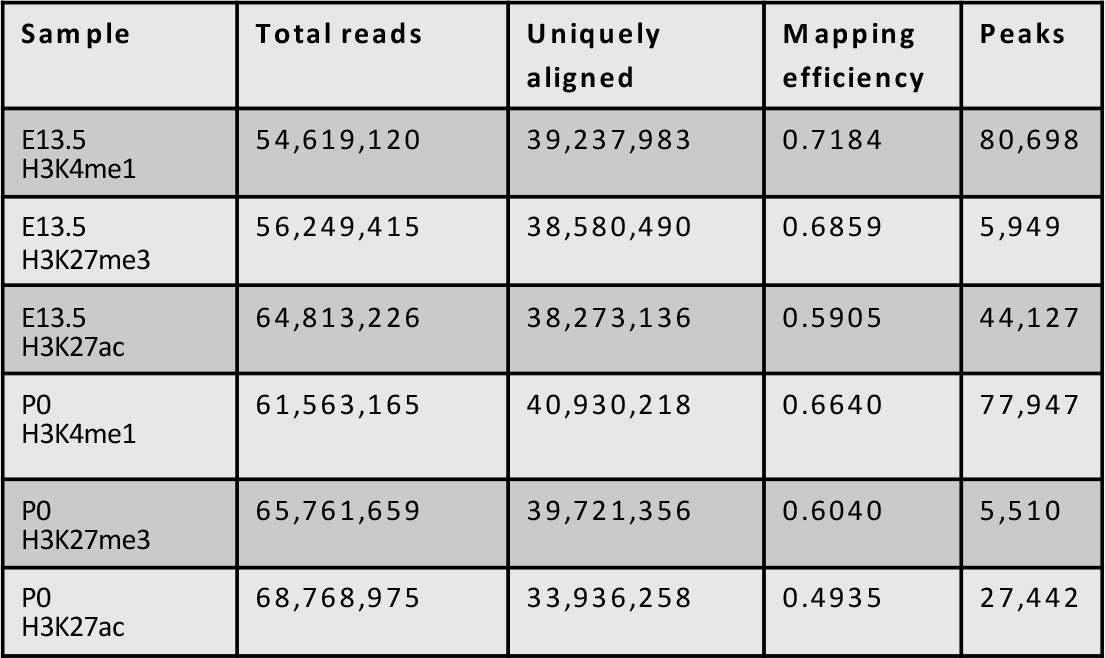
Statistics of ChIP-seq data.

**Supplemental Figure 1.**
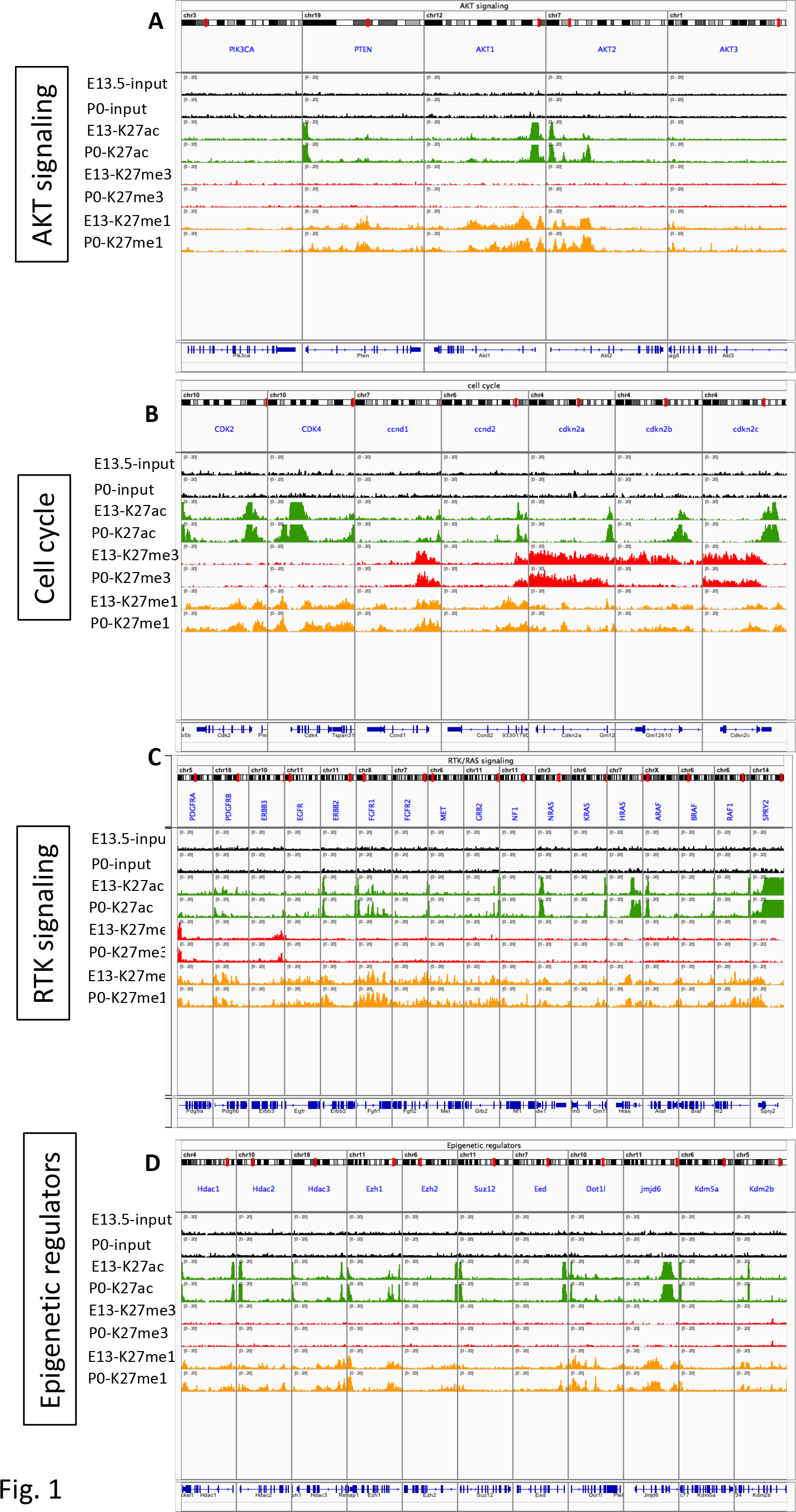
ChIP-seq profiling of histone modifications in representative active pro-renewal genes in young and old NPC.

**Supplemental Figure 2.**
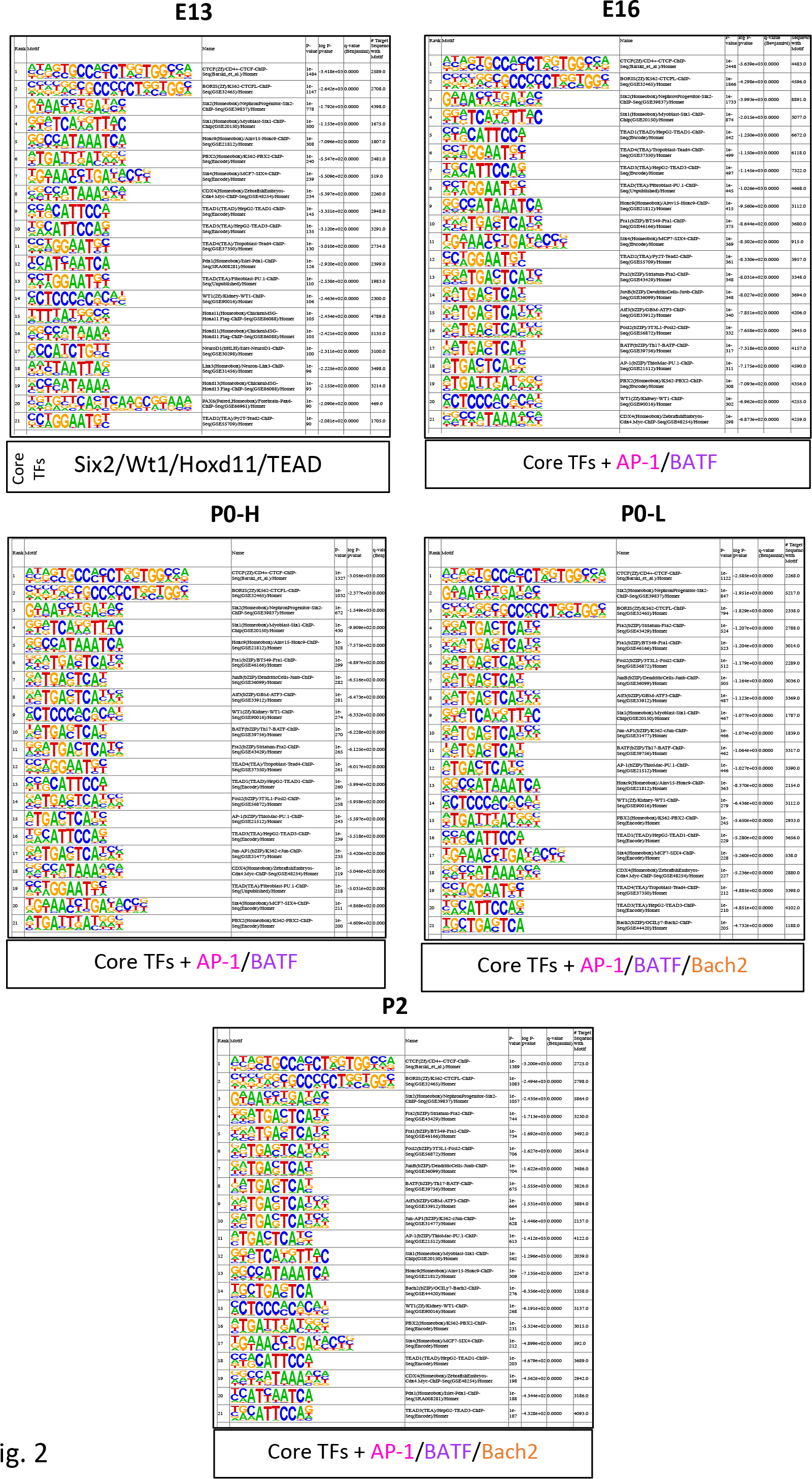
Motif enrichment analysis of ATAC-seq footprints identifies distinct sets of transcription factor binding sites in aging NPC. Binding motifs for the core transcription factors (Six2/Wt1/Hoxd11/Tead) are enriched in NPC of all ages reflecting their shared lineage identity. Open chromatin regions of aging NPC tend to be enriched in binding motifs for AP-1/Bach2/Batf.

